# Individual Variation in Brain Network Topology Predicts Emotional Intelligence

**DOI:** 10.1101/275768

**Authors:** Ling George, Lee Ivy, Guimond Synthia, Lutz Olivia, Tandon Neeraj, Öngür Dost, Eack Shaun, Lewandowski Kathryn, Keshavan Matcheri, Brady Roscoe

## Abstract

**Background:** Social cognitive ability is a significant determinant of functional outcome and deficits in social cognition are a disabling symptom of psychotic disorders. The neurobiological underpinnings of social cognition are not well understood, hampering our ability to ameliorate these deficits.

**Objective:** Using ‘resting-state’ fMRI (functional magnetic resonance imaging) and a trans-diagnostic, data-driven analytic strategy, we sought to identify the brain network basis of emotional intelligence, a key domain of social cognition.

**Methods:** Study participants included 60 participants with a diagnosis of schizophrenia or schizoaffective disorder and 46 healthy comparison participants. All participants underwent a resting-state fMRI scan. Emotional Intelligence was measured using the Mayer-Salovey-Caruso Emotional Intelligence Test (MSCEIT). A connectome-wide analysis of brain connectivity examined how each individual brain voxel’s connectivity correlated with emotional intelligence using multivariate distance matrix regression (MDMR).

**Results:** We identified a region in the left superior parietal lobule (SPL) where individual network topology predicted emotional intelligence. Specifically, the association of this region with the Default Mode Network predicted higher emotional intelligence and association with the Dorsal Attention Network predicted lower emotional intelligence. This correlation was observed in both schizophrenia and healthy comparison participants.

**Conclusion:** Previous studies have demonstrated individual variance in brain network topology but the cognitive or behavioral relevance of these differences was undetermined. We observe that the left SPL, a region of high individual variance at the cytoarchitectonic level, also demonstrates individual variance in its association with large scale brain networks and that network topology predicts emotional intelligence.

## Introduction

Social cognition is a multidimensional construct encompassing a number of mental processes related to perception of, interpretation of, and response to the social environment.(Green et al., 2008). Deficits in multiple domains of social cognition are well-described in patients with psychotic disorders ((Bora et al., 2009; Dodell-Feder et al., 2014; Green et al., 2015; Kohler et al., 2010; Mehta et al., 2014; Savla et al., 2013; Sprong et al., 2007). These deficits may be both partially independent of neurocognitive deficits and are strongly associated with functional status (Allen et al., 2007; Fett et al., 2011; Green, 2016; Hoe et al., 2012; Mehta et al., 2013; Sergi et al., 2007). These findings suggest that social cognitive functioning may be underpinned by neurobiological processes that are at least partially distinct from those associated with other cognitive processes, and contribute uniquely to poor functional outcomes in schizophrenia.

Studies of the neuroanatomical basis of social cognitive ability have frequently utilized functional magnetic resonance imaging (fMRI) in both clinical and non-clinical populations to identify brain regions associated with various domains of social processing (see (Green et al., 2015) for review). Recent studies examining task-free (‘resting state’) brain connectivity have found abnormal resting state connectivity in medial prefrontal and temporal networks that is correlated with social cognitive dysfunction (Abram et al., 2017).. Functional connectivity in these areas, commonly associated with the default mode network (DMN), has been linked to social cognition and real world social functioning in participants with schizophrenia, first-degree relatives of people with schizophrenia and healthy comparison participants (Dodell-Feder et al., 2014; Fox et al., 2017).

One key domain of social cognition is emotional intelligence, ‘the subset of social intelligence that involves the ability to monitor one’s own and others’ feelings and emotions, to discriminate among them and to use this information to guide one’s thinking and actions’ (Salovey and Mayer, 1990). Emotional intelligence is commonly measured using the Mayer-Salovey-Caruso Emotional Intelligence Test (MSCEIT) (Mayer et al., 2003). This test reliably detects social cognitive deficits and is a predictor of social functioning (Eack et al., 2010; Nuechterlein et al., 2008). As such, it was chosen as the measure of social cognition in the battery of tests recommended by the National Institutes of Mental Health Measurement and Treatment Research to Improve Cognition in Schizophrenia (MATRICS)(Marder, 2006).

In this study we sought to identify the brain network correlates of emotional intelligence. We made two methodological decisions in our approach:

First, published hypothesis-driven approaches have been constrained in their ability to capture findings outside the experimental model. We therefore chose to conduct a connectome-wide, entirely data-driven approach to elucidate relationships between social cognition and connectivity.

Second, deficits in social cognitive ability are related to functional outcome in both clinical and non-clinical populations (Dodell-Feder et al., 2014). We sought to discover common dimensional relationships between cognition and connectivity. We hypothesized that the combination of a connectome-wide data analysis and a study sample of schizophrenia and healthy comparison participants would demonstrate trans-diagnostic relationships between connectivity and cognition.

We determined the relationship between emotional intelligence and functional connectivity at the level of individual brain voxels using multivariate distance matrix regression (MDMR), a technique for connectome-wide association studies (Shehzad et al., 2014). We examined this correlation in a group of over one hundred participants recruited across three distinct sites including individuals with schizophrenia or schizoaffective disorder as well as healthy comparison participants.

We hypothesized that this approach would identify brain networks that mediate emotional intelligence trans-diagnostically and that hubs of the DMN would be included in these findings.

## Methods and Methods

### Participants

The study was approved by the Institutional Review Boards of the University of Pittsburgh (Pittsburgh, PA), McLean Hospital (Belmont, MA), and Beth-Israel Deaconess Medical Center (Boston, MA), and all participants gave written informed consent before participating. Participants were recruited from health centers using a variety of means including early psychosis treatment programs and community referral networks. Participants at the Boston and Pittsburgh sites were recruited for a clinical trial (BICEPS, NCT01561859). Clinical and imaging data analyzed here was from participants’ baseline (pre-intervention) evaluation.

Diagnosis was determined using the Structured Clinical Interview for the DSM-IV (SCID) (First and New York State Psychiatric Institute. Biometrics Research, 2007). Patients were assessed by raters trained in the administration and scoring of the SCID. Inclusion criteria for all participants included: (1) age 18-45 years; (2) current IQ ≥ 80 as assessed using the WASI-II ^46^; and (3) the ability to read (sixth grade level or higher) and speak fluent English. Additional inclusion criteria for psychotic disorder participants were (1) a diagnosis of schizophrenia or schizoaffective disorder verified using the SCID interview ^45^; (2) time since first psychotic symptom of ≤ 8 years; (3) clinically stabilized on antipsychotic medication (assessed via SCID in consensus conferences); Healthy comparison (HC) participants did not meet criteria for any Axis I psychiatric disorder currently or historically.

Exclusion criteria were (1) significant neurological or medical disorders that may produce cognitive impairment (e.g., seizure disorder, traumatic brain injury); (2) persistent suicidal or homicidal behavior; (3) a recent history of substance abuse or dependence (within the past 3 months); (4) any MRI contraindications and (5) decisional incapacity requiring a guardian.

Demographic, clinical, and medication regimen information are summarized in Supplemental Table 1.

### Cognitive Testing

Participant emotional intelligence was assessed using the MATRICS Consensus Cognitive Battery (MCCB) (Nuechterlein et al., 2008). This testing battery includes 7 composite scores of processing speed, attention, working memory, verbal learning, visual learning, problem solving, and social cognition. The social cognition score is calculated from the Mayer-Salovey-Caruso Emotional Intelligence Test (MSCEIT) “managing emotions” subscale (Mayer et al., 2003). The MSCEIT includes a series of vignettes. Participants are presented with a series of possible actions related to each vignette and asked to assess the effects of each action on the actor’s or other characters’ mood states or behaviors. Responses are based on a Likert-type scale. We used age and sex normed T scores, which were calculated using the MCCB scoring package.

### MRI data acquisition

Boston & Belmont sites: Data were acquired on 3T Siemens Trio (TIM upgrade) scanners using a standard head coil. The echoplanar imaging parameters were as follows: repetition time, 3000 milliseconds; echo time, 30 milliseconds; flip angle, 85°; 3 × 3 × 3-mm voxels; and 47 axial sections collected with interleaved acquisition and no gap. Structural data included a high-resolution T1 image. In addition, all participants underwent a resting fMRI run with the instructions ‘remain still, stay awake, and keep your eyes open’. Each functional run lasted 6.2 minutes (124 time points).

Pittsburgh site: Data were acquired on a 3T Siemens Verio scanner using a standard head coil. The echoplanar imaging parameters were as follows: repetition time, 3000 milliseconds; echo time, 30 milliseconds; flip angle, 85°; 3 × 3 × 3-mm voxels; and 45 axial sections collected with interleaved acquisition and no gap. Structural data included a high-resolution T1 image. In addition, all participants underwent a resting fMRI run. that lasted 6.2 minutes (124 time points).

### MRI data processing

The imaging data were preprocessed using DPABI image processing software (Yan et al., 2016). To minimize effects of scanner signal stabilization, the first four images were omitted from all analysis. Scans with head motion exceeding 3 mm or 3° of maximum rotation through the resting-state run were discarded. Functional and structural images were co-registered. Structural images were then normalized and segmented into gray, white and CSF partitions using the DARTEL technique(Ashburner, 2007) A Friston 24-parameter model(Friston et al., 1996) was used to regress out head motion effects from the realigned data. CSF and white matter signals, global signal as well as the linear trend were also regressed out. After realigning, slice timing correction, and co-registration, framewise displacement (FD) was calculated for all resting state volumes (Power et al., 2012). All volumes with a FD greater than 0.2mm were regressed out as nuisance covariates. Any scan with 50% of volumes removed was discarded. After nuisance covariate regression, the resultant data were band pass filtered to select low frequency (0.01-0.08Hz) signals. Filtered data were normalized by DARTEL into MNI space and then smoothed by a Gaussian kernel of 8mm^3^ full-width at half maximum (FWHM). Voxels within a group derived gray matter mask were used for further analyses.

After preprocessing, a total of 60 participants with schizophrenia or schizoaffective disorder, and 46 HC participants across all sites remained in the study (Supplemental Table 1).

### Functional Connectivity Analysis

#### Multivariate Distance Matrix Regression

We performed a connectome-wide association study using multivariate distance matrix regression (MDMR) as originally described in (Shehzad et al., 2014). As has been previously described (Satterthwaite et al., 2015; Shanmugan et al., 2016; Sharma et al., 2017) this analysis occurs in three stages: First, a seed-to-voxel connectivity map is generated by using an individual voxel’s BOLD signal timecourse and calculating the Pearson’s correlation coefficients between that voxel and all other gray matter voxels. These maps are generated for every participant. In the second stage, the pattern of 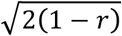 connectivity is compared among participants using the distance metric. In the third stage, multivariate distance matrix regression tests the relationship between inter-subject differences on a given variable (here MSCEIT score) and inter-subject distances on the connectivity maps generated in stage two. This test results in a pseudo-F statistic for each voxel that demonstrates how MSCEIT score is reflected in functional connectivity at that voxel. This process is repeated for every voxel. Scanner site was included as a covariate in this analysis. Type I error was controlled using a voxel height of p<.001 and a cluster extent probability of p<.05 (Figure 1A).

**Figure 1.**
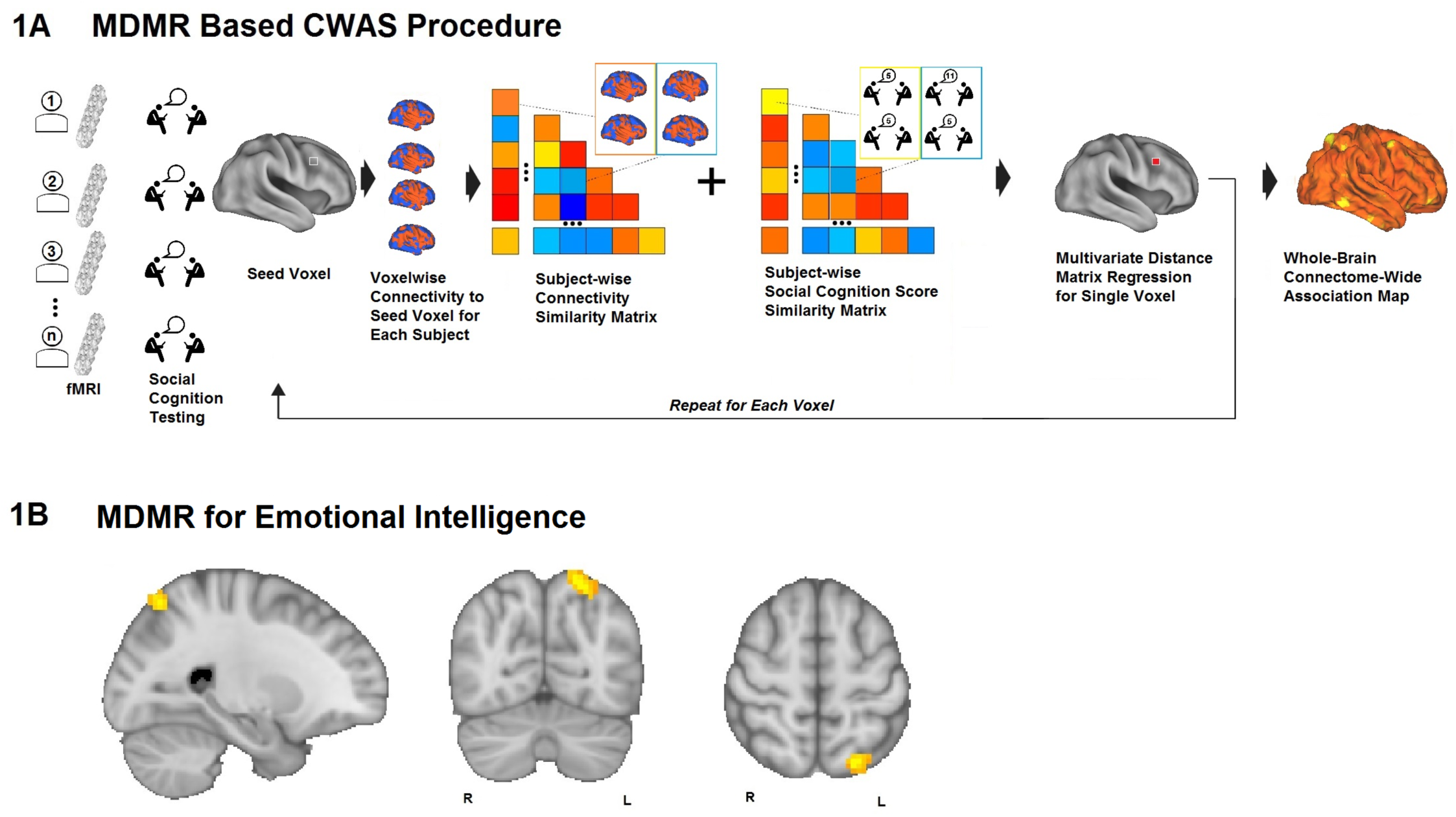
Multivariate Distance Matrix Regression identifies brain voxels whose functional connectivity varies with performance on a measure of emotional intelligence. **1A**. MDMR procedure: rsfMRI and emotional intelligence testing are collected from each participant. For each participant a functional connectivity map is generated to an individual voxel. The pattern of connectivity is compared among participants using a distance metric. Next, multivariate distance matrix regression tests the relationship between inter-subject differences on emotional intelligence and inter-subject distances on the functional connectivity maps. This test results in a pseudo-F statistic for each voxel that demonstrates how emotional intelligence is reflected in functional connectivity at that voxel. This process is repeated for every voxel. **1B**. In our sample of 106 participants (n=60 with schizophrenia or schizoaffective disorder and n= 46 HC participants), we identified a single region in the left superior parietal lobule (extent k=59, centered at MNI coordinates X-24 y-69 z57) whose connectivity correlated significantly with emotional intelligence. Connectivity is thresholded at a voxelwise level of p<.001 and extent threshold of p<.05.

This analysis identifies regions of interest (ROI) where MSCEIT score correlates with functional connectivity. The steps of the analysis described above do not display the pattern or direction (i.e. correlation vs anti-correlation) of connectivity that generated the result. To visualize these patterns, it is necessary to conduct post-hoc testing of the regions of interest identified from MDMR. This ROI-based analysis allows visualization of the connectivity maps that generated the initial result. As others have emphasized (Satterthwaite et al., 2015; Shanmugan et al., 2016; Sharma et al., 2017), these post-hoc tests are not new hypothesis tests, rather, they allow visualization of the connectivity that gave rise to the original MDMR result.

### ROI based analyses

For ROI based analyses we used DPABI to extract the time course of the BOLD signal in a given ROI, then generated whole brain maps of z-transformed Pearson’s correlation coefficients. These maps were entered into SPM12 (Statistical and Parametric Mapping, http://www.fil.ion.ucl.ac.uk/spm). For ROIs generated from MDMR above, we regressed the maps against MSCEIT score to generate maps of how whole brain functional connectivity to the ROIs varies with MSCEIT score. SPM analyses included site as a covariate as in the MDMR analysis. To control for participant demographic variables, we repeated these analyses with age and sex as additional covariates. To control for movement effects we re-performed these analyses with individual mean FD (framewise displacement) as a subject-level correction for motion effects in all subjects (Power et al., 2012). To control for the effects of medication regimen, the analyses were re-performed with prescribed antipsychotic dosage (in chlorpromazine equivalents, CPZE) as a covariate within the schizophrenia/schizoaffective group.

For other, subsequent ROI based connectivity analyses described below, we used DPABI to calculate the time course of the average BOLD signal in a sphere, and then generated whole brain maps of z-transformed Pearson’s correlation coefficients. We then used SPM12 to generate single-sample t-tests of these connectivity maps.

### Structural MRI Analysis

Structural MPRAGE images were checked for artifacts and were processed using Freesurfer 6 (http://surfer.nmr.mgh.harvard.edu/). The images went through first-level auto-reconstruction to register the scans in standard space and skull strip the brains. One rater (OL) edited the images to remove dura, sinuses and vessels that could interfere with segmentation. After visual inspection, 98 images were retained for analysis. The images then went through second and third level auto-reconstruction to register the brains to the Desikan-Killiany atlas and extract sulcal depth measurements (Desikan et al., 2006). Sulcal depth in Freesurfer is the integrated dot product of the movement vector and the surface normal during inflation. Deep regions therefore are expressed as positive values as they move outwards while more superficial regions move inwards. Units are the standard deviation from the median value.

### Table and figure generation

Graphs of relationships between emotional intelligence and structural/functional measures were generated using R. Projections of ROIs and T contrast maps onto cortical surfaces was accomplished using FSL (Jenkinson et al., 2012) and Caret (Van Essen et al., 2001).

## Results

### Functional Connectivity in the Superior Parietal Lobule Predicts Emotional Intelligence

MDMR analysis performed across all 106 participants (60 schizophrenia, 46 HC) revealed a single region whose intrinsic functional connectivity correlated significantly with emotional intelligence. This identified a (59 voxel) region in the left superior parietal lobule (SPL) centered at MNI coordinates X-24 y-69 z57 (Figure 1B).

### SPL Association with Distributed Brain Networks Predicts Emotional Intelligence

We performed Post-hoc testing using this SPL region as a ROI to determine the distribution and directionality of connectivity that gave rise to this result. This analysis revealed two apparent patterns of functional connectivity: Higher emotional intelligence correlated with increasing intrinsic connectivity between the left SPL and bilateral fronto-parietal and temporal regions that correspond to the canonical “default mode network” (DMN) (Figure 2A). Inversely, lower emotional intelligence correlated with increasing functional connectivity between the left SPL and bilateral parietal-occipital regions that mirror the canonical distribution of the “dorsal attention network” (DAN) (Figure 2B). This result remained significant with the inclusion of sex and age as covariates and was significant with individual motion (FD) included as a covariate.

**Figure 2.**
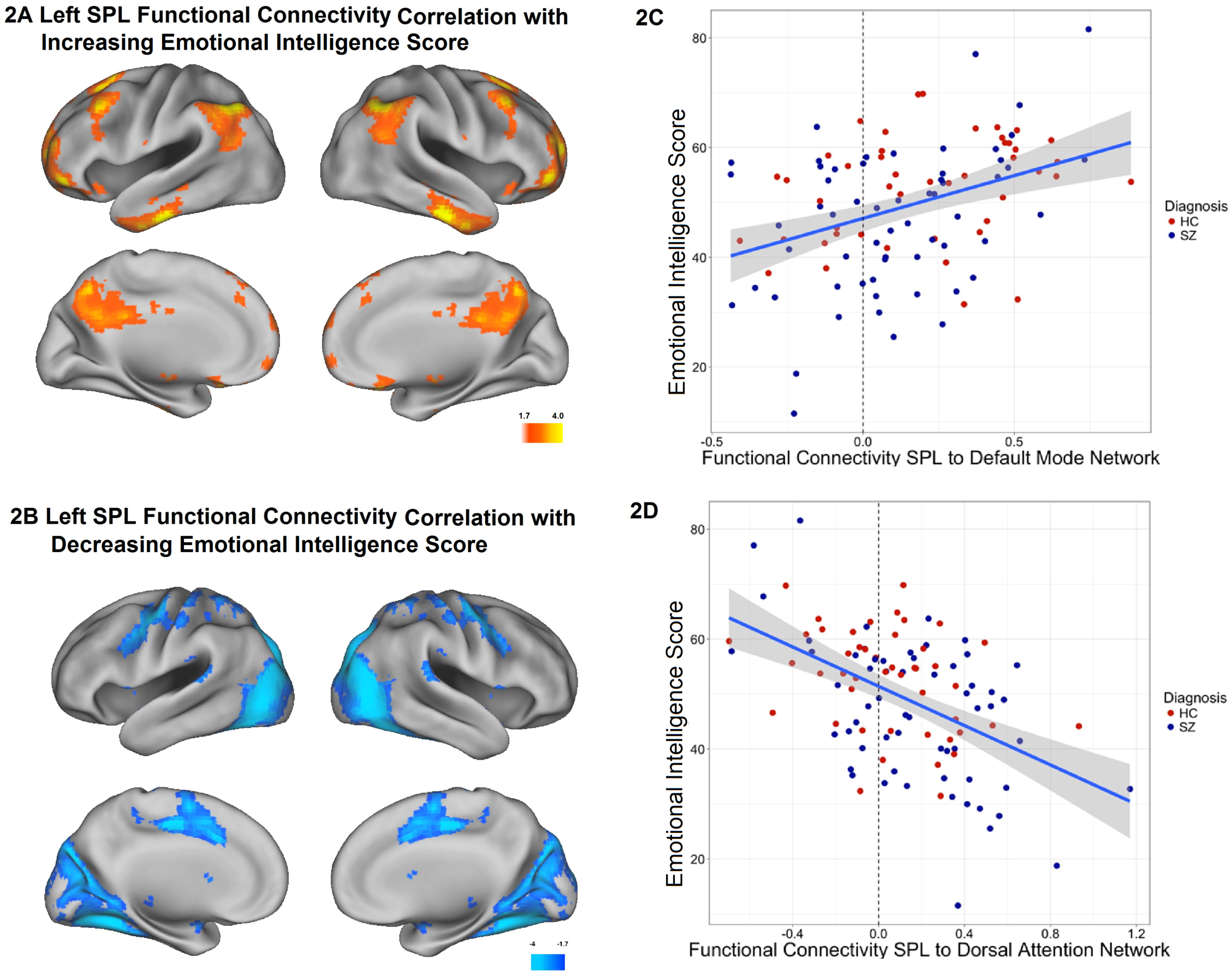
Post-hoc testing to determine the pattern and directionality of connectivity that gave rise to the MDMR result. Two patterns of connectivity are observed. **2A**. Emotional intelligence is correlated with functional connectivity between the left SPL region and the “default mode network” (DMN). **2B**. Emotional intelligence is inversely correlated with functional connectivity between the left SPL and the “dorsal attention network” (DAN). Color bar: T-statistic. **2C**. Plot of emotional intelligence score (y-axis) and functional connectivity between 6mm sphere at SPL ROI coordinates (X-24 y-69 z57) and 6mm sphere at the region of maximally correlated connectivity in the DMN (x12 y-87 z-30) (x-axis). **2D**. Plot of emotional intelligence (y-axis) and functional connectivity between 6mm sphere at SPL ROI coordinates X-24 y-69 z57 and 6mm sphere at the region of maximal inversely correlated connectivity in the DAN (x36 y-78 z21) (x-axis). Notably, both graphs demonstrate correlations that cross the reference line of null functional connectivity, meaning that this ROI does not simply display cognition-connectivity correlations within the DMN and expected cognition-connectivity anti-correlations in the network (DAN) anti-correlated with the DMN. Rather, the network association of this region appears to shift from being a member of the DMN to a member of the DAN as one moves along the distribution from higher scoring individuals to lower scoring individuals.

We plotted the relationship between emotional intelligence and functional connectivity for both networks as a scatter plot in Figure 2C & 2D. As expected from the maps in Figure 2A & 2B, we observed that higher emotional intelligence was correlated with increased functional connectivity between the SPL and the DMN and decreased connectivity to the DAN. Because the DMN is intrinsically anti-correlated with the DAN, this result could be expected for any region in one of these networks (i.e. a region of interest located in the distribution of the DMN that demonstrated lower correlation to the rest of the DMN would be expected to demonstrate higher correlation to the DAN). Surprisingly, the scatter plots of connectivity cross the axis of 0 connectivity i.e. *this SPL region is intrinsically connected to the DMN in some individuals and is intrinsically connected to the DAN in others*.

### Connectivity Associations with Emotional Intelligence Within Diagnostic Groups

We examined functional connectivity-emotional intelligence relationships in HC participants and those with schizophrenia/schizoaffective disorder independently. Using the same ROIs as in figure 2C & 2D, we correlated FC with emotional intelligence. We observe similar strengths of correlation between emotional intelligence and DMN-SPL connectivity in HC (R=.329, p=.026) and schizophrenia (R= .377, p=.003) samples and also see similar correlation strengths in DAN-SPL connectivity in HC (R=-.445, p=.002) and schizophrenia samples (R=-.504, p<.001).

### Site Replicability of Connectivity-Cognition Relationships

Given significant concerns about the replicability of fMRI imaging studies, we sought to determine if these findings in all of our participants could be observed in each study population (BICEPS and McLean Hospital) independently to better confirm the replicability of this result. Both study populations independently observe the same pattern of SPL functional connectivity correlating with emotional intelligence (i.e. DMN connectivity with higher emotional intelligence and DAN connectivity with lower emotional intelligence) (Supplemental Figure 1).

### Domain Specificity of SPL Connectivity-Cognition Relationships

As an exploratory analysis, we asked if the network connectivity of the SPL region predicted cognition broadly or was specific to emotional intelligence. We examined connectivity to this ROI regressed against MSCEIT score with full scale IQ (FSIQ) as a covariate. This was performed on participants in the study with FSIQ data available (BICEPS i.e. Pittsburgh and Boston). In this analysis, we again observed the same pattern of emotional intelligence correlated functional connectivity as seen without FSIQ as a covariate (Supplementary Figure 2).

### Network Toploogy Predicts Emotional Intelligence

In order to investigate the relationship between cognition and inter-individual variance in network topography, we performed an exploratory post-hoc analysis to visualize intrinsic connectivity to this region in subsets of participants. We performed one-sample T-tests among the participants with the highest and lowest emotional intelligence. In this analysis, we examined intrinsic SPL connectivity among participants top 5^th^ percentile of emotional intelligence and participants in the bottom 5^th^ percentile of emotional intelligence. Using this analysis, the SPL ROI in the top 5^th^ percentile scorers showed connectivity with the canonical DMN. In the bottom 5^th^ percentile, this ROI was a member of the DAN (figure 3A).

**Figure 3.**
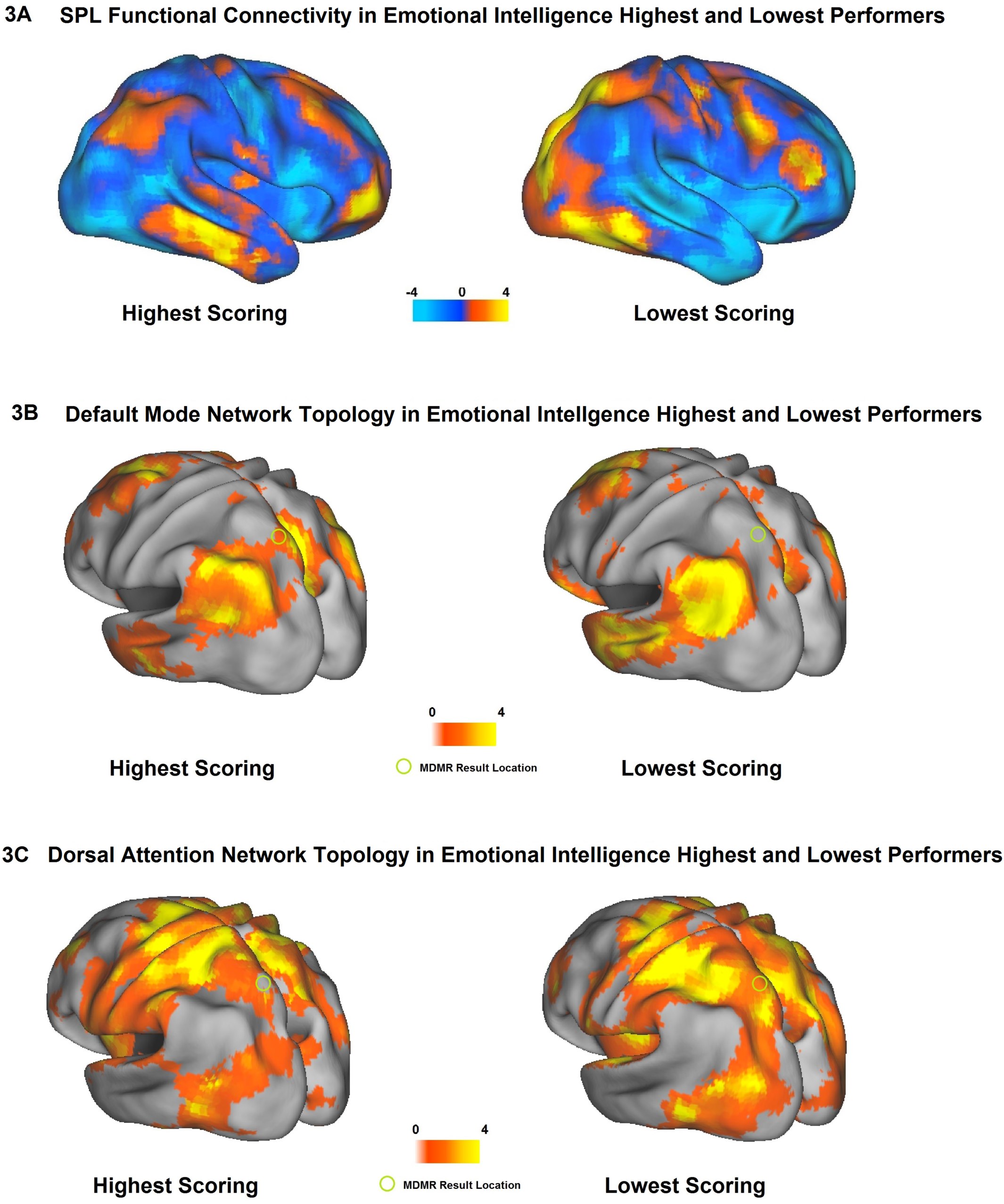
The left SPL ROI is connected to different brain networks in participants with high versus low emotional intelligence. Here we visually confirm the implication of the cognition-connectivity correlations shown in 2C and 2D by looking at absolute connectivity in the highest and lowest scorers on emotional intelligence testing. **3A** One sample T-tests of functional connectivity to the Left SPL ROI in the participants in the top 5^th^ percentile of emotional intelligence and the participants in the bottom 5^th^ percentile. In the highest scoring individuals, the SPL ROI is correlated with the DMN and anti-correlated with the DAN. In the lowest scoring individuals, the SPL ROI is correlated with DAN and anti-correlated with DMN. Color bar = T-statistic. **3B & 3C** The topographic distribution of the Default Mode Network and Dorsal Attention Network as visualized using canonical methods: The topographic distribution of the DMN is derived from functional connectivity to a 10mm spherical ROI placed in the precuneus (x-4, y-58, z44) and the DAN from a 10mm sphere ROI placed in the frontal eye fields (x28 y-8, z52). Consistent with the results of 3A above, we observe significant variability in the topology of both networks with their inclusion or exclusion of the SPL ROI predicting emotional intelligence.

**Figure 4.**
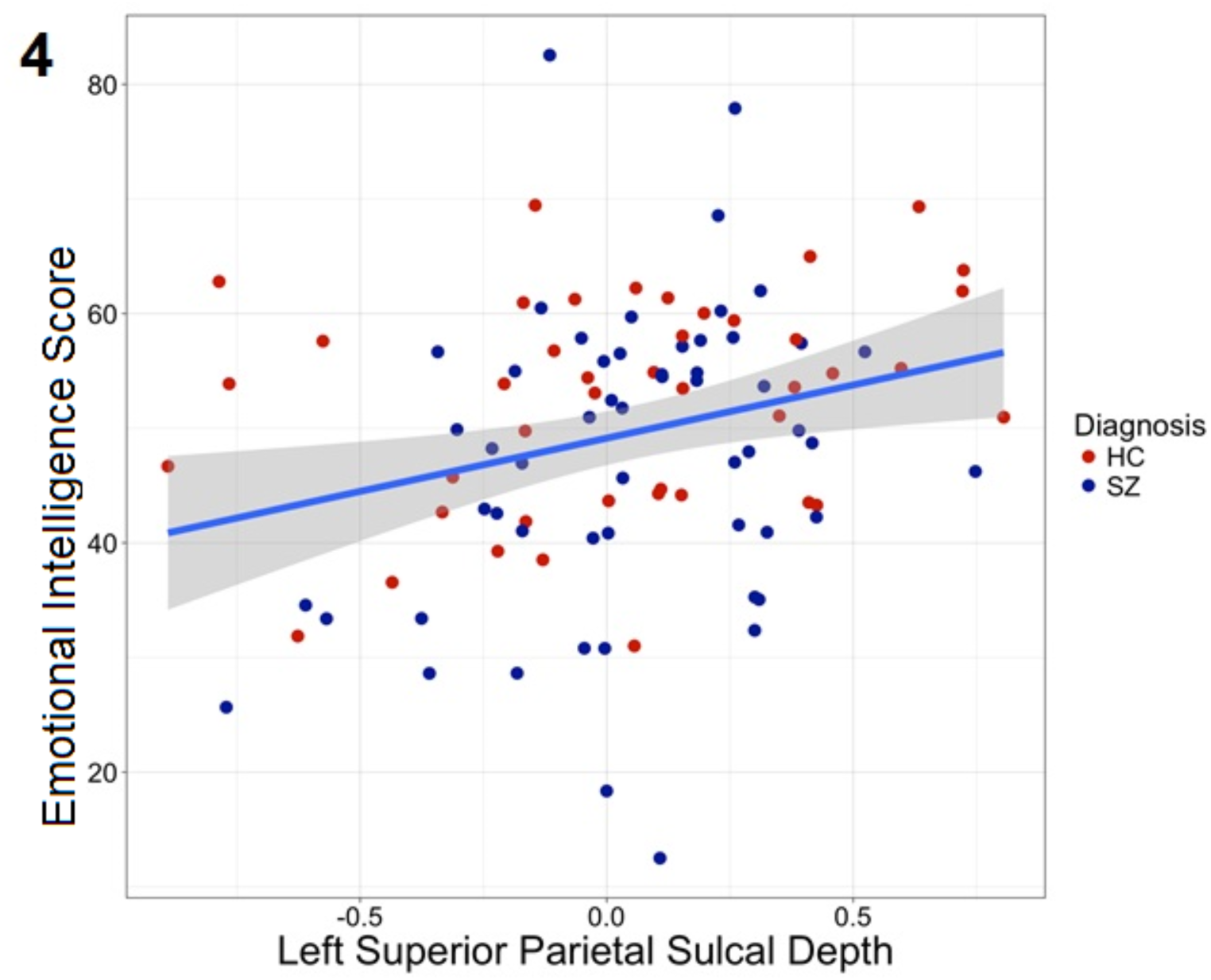
Emotional intelligence is correlated with sulcal depth in the left SPL. The Freesurfer program was used to conduct a ROI based analysis of sulcal depth in the left superior parietal lobule. These values were then correlated with emotional intelligence scores. Units are standard deviations from the median. This analysis revealed a significant correlation between sulcal depth and emotional intelligence (r=0.269, *p*=0.0075).

To better place this finding in the context of existing studies of the DMN, we compared the topology of the DMN in the individuals with the highest emotional intelligence and those with the lowest using canonical methods for identifying the DMN (figure 3B). We placed a 10mm spherical ROI in the precuneus to visualize the distribution of the DMN and a 10 mm spherical ROI in the frontal eye fields to visualize the DAN. As expected from the results in figures 2 & 3A, we observe that, in the highest scoring individuals, the SPL ROI is included in the topology of their DMN and in the lowest scoring individuals, the SPL ROI is included in the DAN.

### Sulcal Depth Associations with Emotional Intelligence

Prior studies have revealed considerable individual variation at the cytoarchitectonic level in this region (see discussion). This level of analysis is not available but we asked whether there is a structural MRI correlate to network connectivity and emotional intelligence in this region. Based on prior studies of structural variability in this region (Mueller et al., 2013), and the suggestion that the intra-parietal sulcus area hIP3 may be associated with a DMN-like pattern of connectivity (Wang et al., 2015b), we hypothesized that emotional intelligence would be correlated with sulcal depth in this region. As predicted, we observed a significant (r=0.269, *p*=0.0075) relationship between sulcal depth and emotional intelligence (Figure 4).

## Conclusions

We here identify a region of the left superior parietal lobule whose intrinsic functionally connectivity correlates significantly with emotional intelligence. Surprisingly, the relationship between connectivity and cognition appears to be mediated by topography.. Specifically, this region is variably connected to two large scale brain networks, the DMN and the DAN. The brain is organized into distributed networks whose topology are typically visualized using group averaging (Power et al., 2011; Yeo et al., 2011). Individual variation in network topology among healthy control participants has been demonstrated previously(Gordon et al., 2017a; Gordon et al., 2017b; Laumann et al., 2015; Mueller et al., 2013; Poldrack et al., 2015; Wang et al., 2015a). To date however, this variability in network topology has not been associated with behavioral or cognitive phenotypes. In our study population, emotional intelligence is predicted by brain network membership: In high performers, spontaneous activity in this region is correlated with the default mode network and anti-correlated with the dorsal attention network. This network membership is reversed in low performers.

What is the basis of this variability in network membership of the SPL? A convergence of results from multiple research methods have demonstrated that the SPL can be subdivided into at least four distinct subregions that are differentiable on the basis of cytoarchitectonic description (Scheperjans et al., 2008), white matter tractography (Mars et al., 2011), and both task evoked and resting state functional connectivity (Wang et al., 2015b). Particularly relevant to this study is the finding that postmortem cytoarchitectonic studies identified considerable individual heterogeneity in the SPL (Scheperjans et al., 2008). The topographical distribution of these subregions in the left SPL was observed to show significant inter-subject variability, particularly in the region referred to as area 7A whose center of gravity corresponds to the region identified in this study.

Using group-averaged functional connectivity maps of the SPL Wang et al. observed specific patterns of resting state connectivity to left SPL subregions (Wang et al., 2015b). That study identified five different “clusters” of resting state connectivity to the left SPL. Notably, the adjacent “L3 and L4” regions in that manuscript demonstrate specific connectivity patterns similar to the DMN and DAN patterns we observe among emotional intelligence high and low performers respectively. These regions appear to correspond to the hIP3 & 7A regions respectively, identified by Scheperjans et al. (Scheperjans et al., 2008). Moving outside of these regions, when we examine connectivity at coordinates that correspond to the other SPL subdivisions i.e. the Scheperjans 5L / Wang L2 region rostral to the MDMR ROI or the Scheperjans 7P / Wang L5 caudal to the MDMR ROI, we do not see a correlation between connectivity and emotional intelligence (Supplemental Materials text and Supplemental Figure 3).

We hypothesize that the individual variability in resting state connectivity in this ROI identified by MDMR represents individual variability in the cytoarchitectonic organization of the left SPL. Furthermore, we hypothesize that the correlation between network connectivity and emotional intelligence is a cognitive correlate of this anatomic variability. To date, studies have not linked cytoarchitectonic variation in the SPL to cognitive variation. In support of this hypothesis, one existing study has correlated diffusion tensor imaging measures with performance on the same emotional intelligence test used in this study. In that article, Pisner et al. observed emotional intelligence correlating with increased fractional anisotropy in the cingulum connecting frontal and parietal regions in a distribution entirely consistent with our observation of emotional intelligence correlating with fronto-parietal connectivity (Pisner et al., 2017).

There is converging evidence suggesting that the DMN has a particular role in social cognition (Dodell-Feder et al., 2014; Fox et al., 2017; Mars et al., 2012). The DMN is a large, distributed brain network. Why does such a circumscribed region of this network in the superior parietal lobule appear to be such a critical determinant of emotional intelligence? A neuroanatomical model called the parieto-frontal integration theory (P-FIT) suggests that connectivity between parietal cortex and frontal areas is a critical determinant of intelligence(Jung and Haier, 2007). In this specific scenario, connectivity between the parietal lobule and a parieto-frontal network implicated in social cognition (the DMN) may mediate emotional intelligence..

In our study, the relationship between SPL network membership and emotional intelligence was trans-diagnostic and is observed in both participants with schizophrenia and schizoaffective disorder as well as HC participants. This is observed despite a representative patient sample in which the mean emotional intelligence score was significantly less than that of HC participants (*p*=.015). We hypothesize that what is already known about schizophrenia network pathophysiology may explain social cognition deficits in this disorder. There is evidence that the molecular pathophysiology believed to underlie schizophrenia leads to the selective disruption of fronto-parietal networks (e.g. the DMN) that is a hallmark feature of schizophrenia and related disorders (Baker et al., 2014; Yang et al., 2016). If social cognitive ability is partially dependent on left SPL connectivity to an intact DMN, our result may provide a mechanistic bridge linking a “synaptic” disease to poor societal functioning.

Limitations of this study include the fact the emotional intelligence measure included in the MCCB is one of several sub-tests of the complete MSCEIT test. That said, performance on the four subtests are well described by a single factor solution (Eack et al., 2010). In addition, as with other cohort imaging studies, this study is inherently correlational and cannot distinguish between connectivity directly mediating cognition versus other interpretations e.g. the result of an underlying process that mediates social cognitive ability and network connectivity via unrelated processes.

Ultimately, interventions designed to improve emotional intelligence that incorporate imaging will validate or refute this network-based model of emotional intelligence by measuring network connectivity longitudinally before and after interventions to improve performance. This may be particularly relevant for behavioral clinical trials that utilize a “target engagement” framework to validate experimental therapeutics (Lewandowski et al., 2017). In that vein, the location of our result provides a potential target for neuromodulation. For example, If the network “membership” of this region can be modified via induction of neuroplasticity at this region (e.g. by repetitive transcranial magnetic stimulation), then neuromodulation may enhance trials using cognitive remediation interventions.

More generally, the utility of this result extends beyond imaging studies of social cognition. The demonstration that individual brain network topology is associated with individual variation in complex cognition reinforces a growing appreciation of the limitations of group-based averaging in resting-state imaging. Averaging across groups of subjects likely obfuscates relationships between individual network topology and cognitive and behavioral phenotypes. In this example, the relationship between network topology and cognitive measures was significant enough to be discerned in a group of a hundred subjects scanned a single time each. It seems inevitable that other relationships between network topology and cognition / behavior remain to be discovered but discerning those relationships may likely require “dense” scanning of individual subjects i.e. scanning the same individuals five (Mueller et al., 2013), twelve(Gordon et al., 2017b), or twenty-four times (Braga and Buckner, 2017). Such efforts are resource intensive, but if such deep characterization yields greater understanding of network-behavior relationships, it is reasonable to imagine a re-allocation of resources away from participant numbers and towards precise network mapping. A remaining challenge to be addressed is how such a change in approach may be applied to better understand the network basis of neurological and psychiatric disorders where acquiring multiple scans in symptomatic individuals may prove difficult.

## Supplementary Materials

**Supplemental Table 1.**
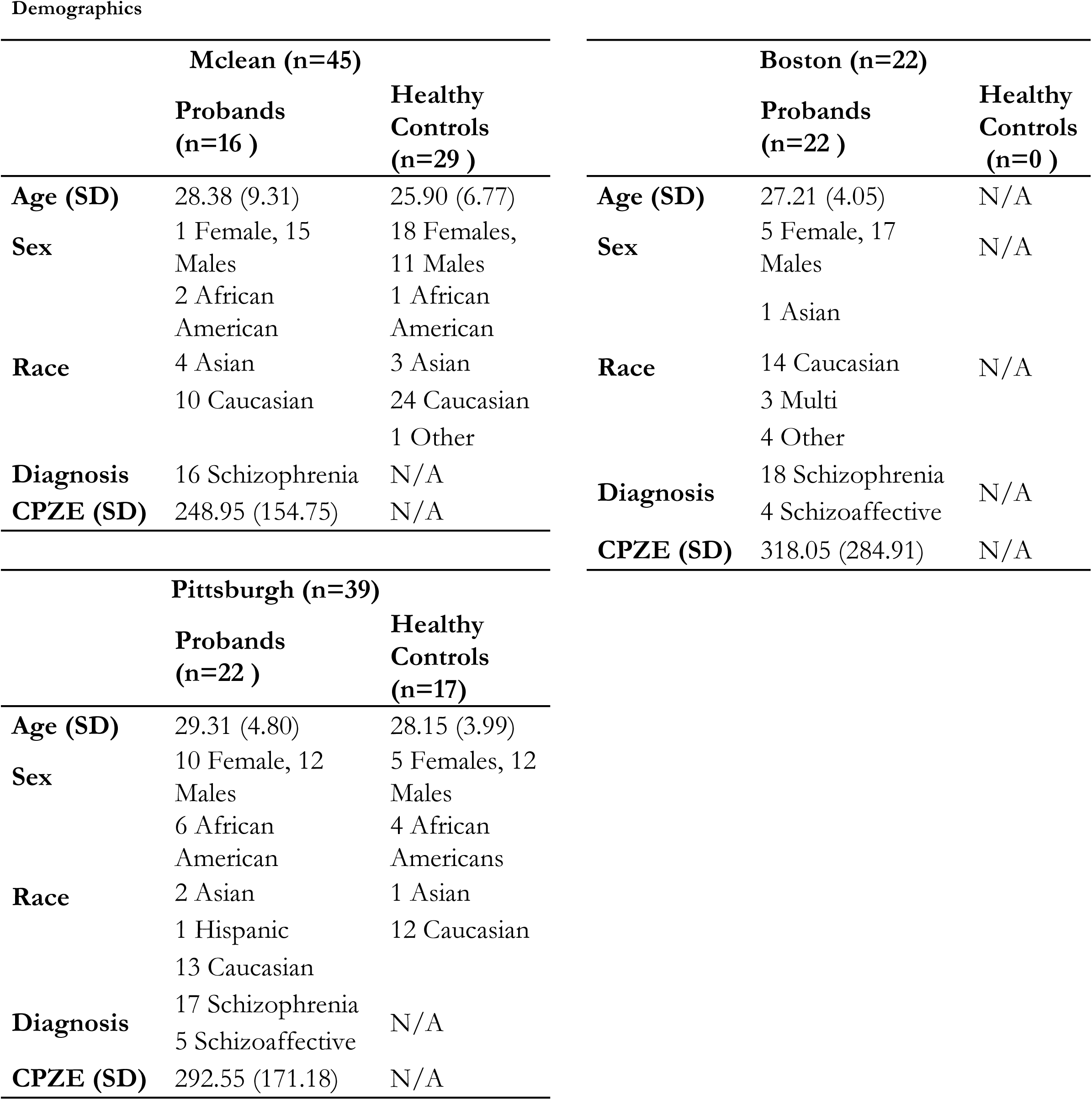
Participant demographics and clinical information. Abbreviations: CPZE chlorpromazine equivalents SD Standard Deviation

**Supplemental Figure 1.**
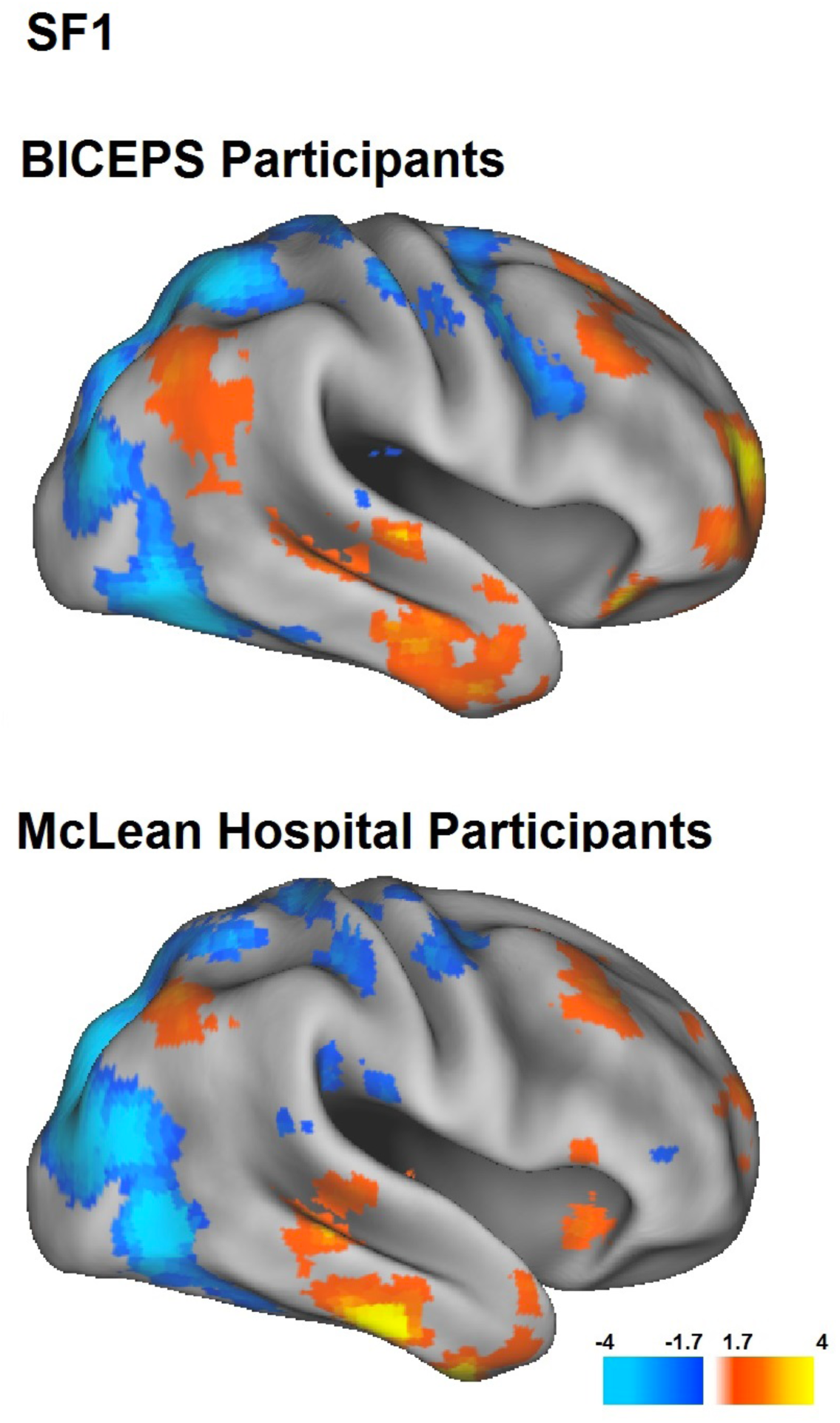
Post-hoc testing of emotional intelligence / functional connectivity correlations in each study population. The same pattern of correlations between left SPL connectivity and emotional intelligence are observed at the McLean and Boston/Pittsburgh (BICEPS) samples independently. Color bar = T-statistic.

**Supplemental Figure 2.**
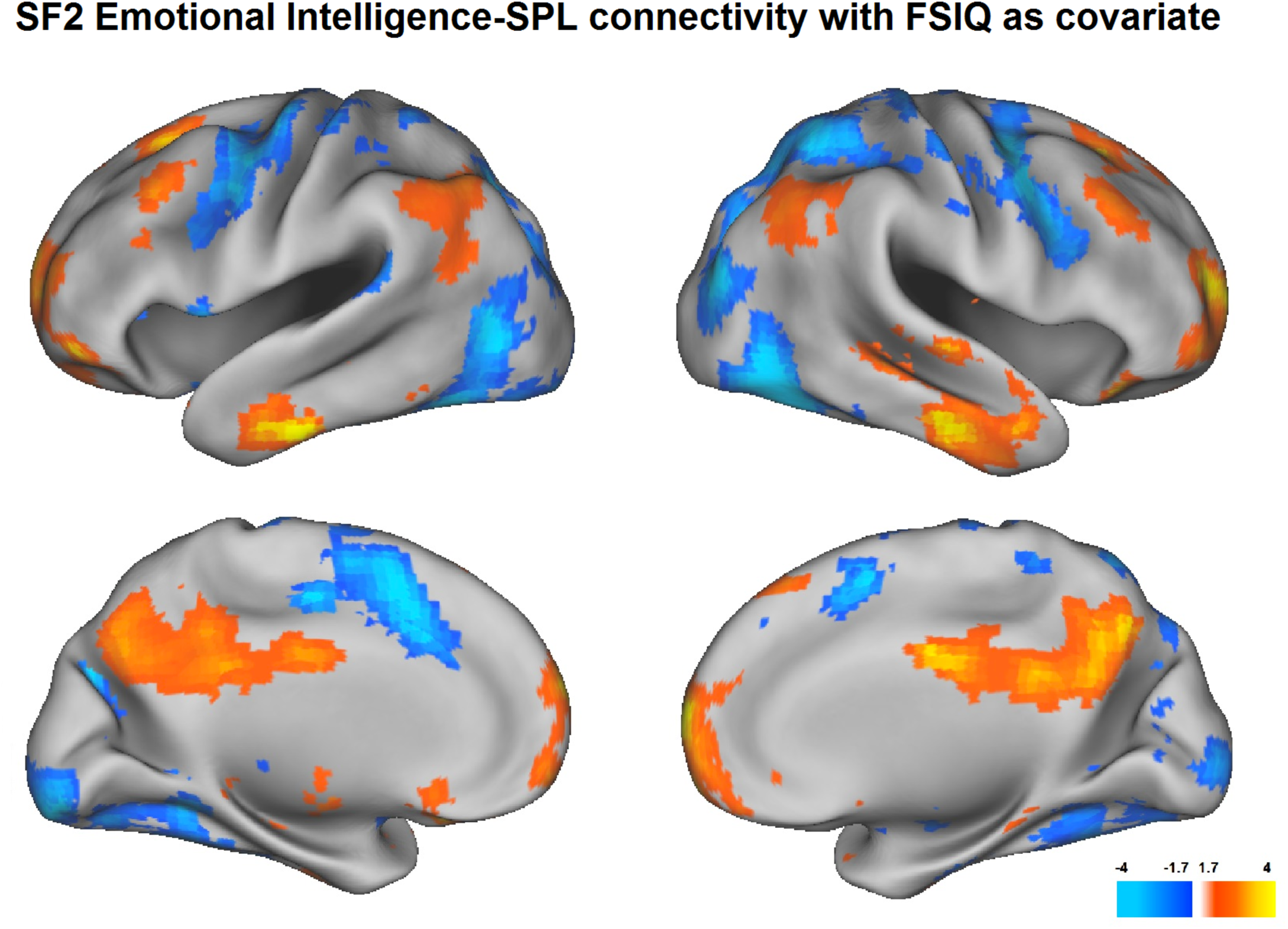
The functional connectivity of the SPL region regressed against emotional intelligence with full scale IQ (FSIQ) as a covariate. In an analysis of the sites with FSIQ available (Pittsburgh and Boston), we continue to observe the same pattern of SPL connectivity-emotional intelligence correlations. Color bar = T-statistic.

### The relationship between functional connectivity and emotional intelligence is SPL subdivision specific

MDMR identifies regions whose functional connectivity correlates significantly with a given variable, in this case emotional intelligence. Adjacent SPL subdivisions were not identified as regions with significant correlation between emotional intelligence and functional connectivity. To better visualize these negative results in adjacent SPL subdivisions, we conducted one-sample connectivity analyses to ROIs in SPL subdivisions just rostral and caudal to the region identified by MDMR. We placed 6mm spherical ROIs at coordinates that correspond to adjacent but cytoarchitectonically distinct regions of the parietal lobe (rostral MNI coordinates x-21 y-55 z72, corresponding to Scheperjans 5L / Wang L2 and caudal MNI coordinates x-24 y-69 z57, corresponding to Scheperjans 7P / Wang L5 (Scheperjans et al., 2008; Wang et al., 2015b).

We examined the intrinsic connectivity of these ROIs and compared them to the ROI identified by MDMR. While participants with high emotional intelligence were readily distinguished from low performers on the basis of connectivity to the MDMR ROI, in the adjacent SPL subdivisions both groups demonstrated largely overlapping connectivity in patterns consistent with previously published results (Wang et al., 2015b) (Supplemental Figure 3). Only the ROI corresponding to the MDMR result demonstrated a relationship between emotional intelligence and network “membership” i.e. connectivity to DMN versus DAN.

**Supplemental Figure 3.**
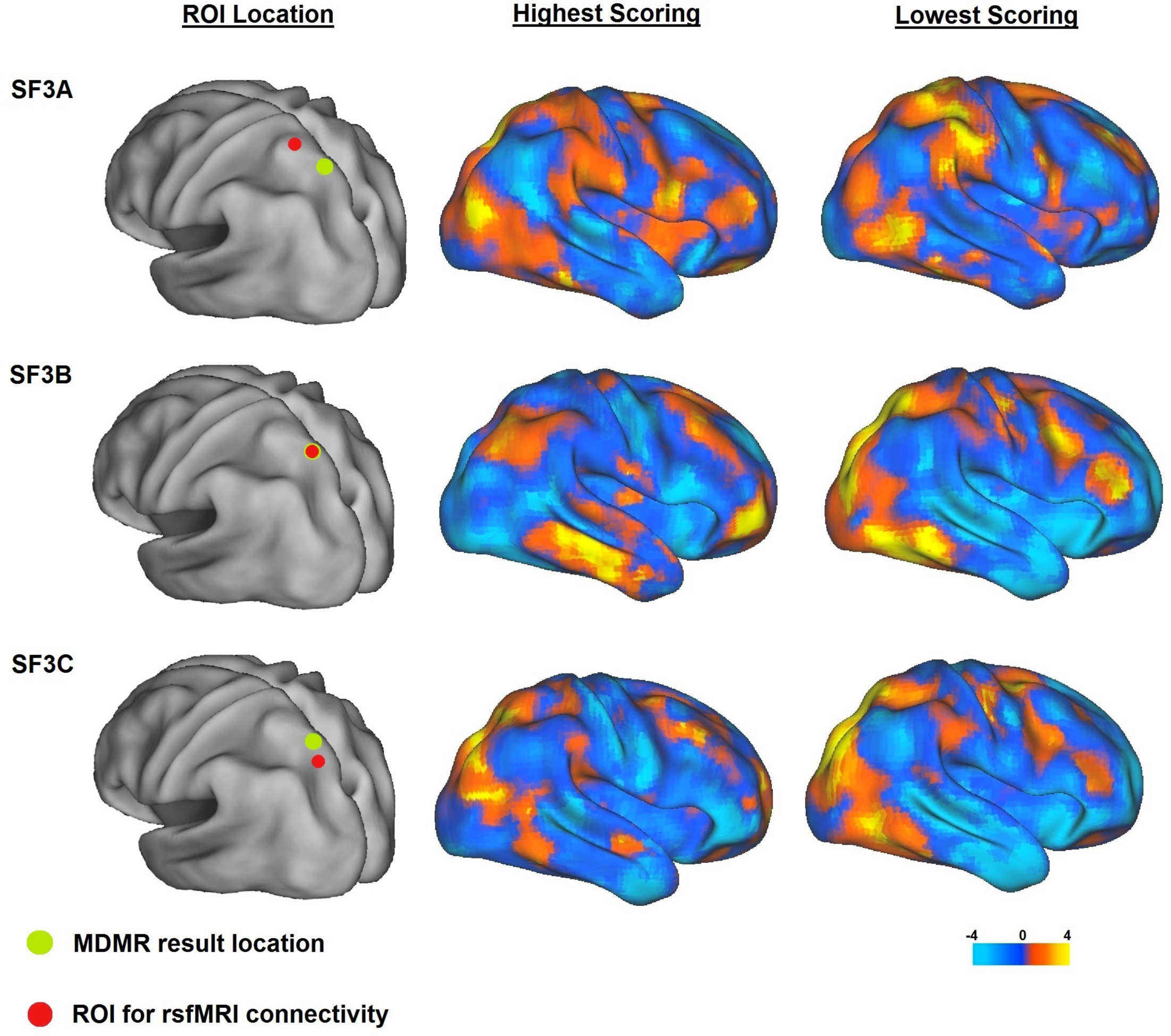
The relationship between functional connectivity and emotional intelligence is SPL subdivision specific. One sample T-tests of functional connectivity to three adjacent superior parietal ROIs in the participants in the top 5^th^ percentile of emotional intelligence and the participants in the bottom 5^th^ percentile of emotional intelligence. These three ROIs are: **SF3A**. 6mm sphere rostral to the MDMR result (SPL region 5L/L2, MNI x-21 y-55 z72). **SF3B**. The MDMR result (MNI x-24 y-69 z57), and **SF3C**. 6mm sphere caudal to the MDMR result (at SPL region 7P/L5, MNI x-16 y-75 z50). Functional connectivity to the MDMR result ROI shows distinct patterns of functional connectivity in the highest and lowest performers in emotional intelligence. The rostral and caudal ROIs demonstrate patterns of connectivity that are largely overlapping between highest and lowest performers on emotional intelligence testing.

## Funding

This work was supported by the National Institutes of Health:

K23MH100623 (RB)

RO1MH92440 (SE, MK)

K24MH104449 (DO)

K23MH091210 (KL)

## Conflicts of interest

DO: Served on Scientific Advisory Board for Neurocrine Inc in 2016 All other authors report no conflicts of interest

This work is not peer-reviewed.

